# Simultaneous Binding of the N- and C-terminal Cytoplasmic Domains of Aquaporin 4 to Calmodulin May Contribute to Vesicular Trafficking

**DOI:** 10.1101/2021.06.29.450403

**Authors:** Hiroaki Ishida, Hans J. Vogel, Alex C. Conner, Philip Kitchen, Roslyn M. Bill, Justin A. MacDonald

## Abstract

Aquaporin 4 (AQP4) is a water transporting, transmembrane channel protein that has important regulatory roles in maintaining cellular water homeostasis. Several other AQP proteins exhibit calmodulin (CaM)-binding properties, and CaM has recently been implicated in the cell surface localization of AQP4 that occurs in response to osmotically-driven changes in cell swelling in the central nervous system. The objective of the present study was to assess the CaM-binding properties of AQP4 in detail. Inspection of AQP4 revealed two putative CaM-binding domains (CBDs) in the cytoplasmic N- and C-terminal regions, respectively. The Ca^2+^-dependent CaM-binding properties of synthetic and recombinant AQP4 CBD peptides were assessed using fluorescence spectroscopy, isothermal titration calorimetry, and two-dimensional ^1^H, ^15^N-HSQC NMR with ^15^N-labeled CaM. The N-terminal CBD peptide of AQP4 predominantly interacted with the N-lobe of CaM with a 1:1 binding ratio and a K_d_ of 3.4 μM. CaM bound two C-terminal AQP4 peptides with interactions observed for both the C- and N-lobes of CaM (K_d1_: 3.6 μM, K_d2_: 113.6 μM, respectively). A recombinant AQP4 protein domain (rAQP4ct, containing the entire cytosolic C-terminal domain sequence) bound CaM in a 1:1 binding mode with a K_d_ of 6.1 μM. A ternary bridging complex could be generated with the N- and C-lobes of CaM interacting simultaneously with the N- and C-terminal CBD peptides. These data suggest that this unique adapter protein binding mode of CaM and AQP4 may be an important regulatory mechanism for the vesicular trafficking of AQP4.

## 1. INTRODUCTION

Calmodulin (CaM) is a small eukaryotic protein of approximately 17 kDa that functions as the primary receptor of intracellular [Ca^2+^] to elicit the regulation of a diverse group of cellular proteins [1-4]. CaM binds four Ca^2+^ ions using four EF-hand binding sites that are equally divided between two globular domains called the N-lobe and the C-lobe [5]. The binding of Ca^2+^ leads to conformational changes that enable Ca^2+^-CaM to interact with many different target proteins. The calmodulin-binding domains (CBDs) that are present in target proteins do not display a strict consensus sequence; however, many targets bind CaM in a similar “canonical” manner [1, 2, 4, 6-8]. In this case, Ca^2+^-CaM interacts with its target using both the N- and C-lobes in a 1:1 ratio. CBD sequence motifs that provide canonical CaM-binding usually provide an amphiphilic α-helical structure, generate a net positive charge in the binding region, and contain at least two hydrophobic anchor residues near the ends of the sequence. The positioning and distance between the anchor residues are thought to define the binding mode of the CBD and its orientation with respect to Ca^2+^-CaM. CBDs are generally classified into motifs based on spacing between hydrophobic anchor residues, including 1-8-14, 1-5-10, 1-10, 1-14, and 1-16 among others [2, 4, 7, 8]. The number of protein targets possessing functional CBD motifs continues to grow as experimental data unambiguously reveal their ability to bind CaM [9]. However, not all CaM-complexes follow the canonical 1:1 binding mode, and many complexes with unique interaction motifs have been characterized (for recent review see [1]). In several cases, CaM has been found to act as an adaptor protein that can bridge between two different domains or target proteins [1].

Calmodulin is commonly associated with membrane-spanning channels, and it is sometimes even considered to be an integral subunit of various channel protein complexes [6]. Indeed, CBD motifs and CaM-binding properties of aquaporin (AQP) water channels have previously been reported. The AQPs are a family of tetrameric, integral membrane channels that govern the transport of small solutes (water and in some cases small neutral solutes such as glycerol) across cellular membranes [10-13]. They play important roles in the regulation of body fluid homeostasis in many organ systems. To date, three AQP family members have been reported to interact with CaM, namely AQP0, AQP4 and AQP6. AQP0 channel activity is directly regulated by Ca^2+^ [14, 15], and CaM antagonists could attenuate Ca^2+^-dependent effects on its water permeability [14]. Additional detailed biochemical studies provided evidence for direct CaM-AQP0 interactions [16], and the CaM-binding events were described as associations of Ca^2+^-CaM to a non-canonical CBD located in the intracellular C-terminal helix of AQP0 [17-20]. The binding of CaM to AQP0 could be inhibited by phosphorylation of Ser-235, a residue located within the CBD [16, 18, 20]. Ultimately, a “bind and capture” mechanism for Ca^2+^-CaM regulation of AQP0 channel activity was proposed [20]. It was suggested that the binding of CaM to the C-terminal CBD of two neighboring AQP0 subunits within the tetramer results in stabilization of a closed pore state and restricts channel permeability [19-21]. Less is known about the functional impact of CaM binding to AQP6; however, a CBD motif was tentatively identified within its intracellular N-terminal helix [22]. Finally, recent reports have implicated CaM in the cell surface localization of AQP4 that occurs in response to osmotically-driven changes in the central nervous system, leading to cytotoxic edema [23-25]. This process involves the transfer of AQP4 from intracellular storage vesicles to the plasma membrane. In this regard, the accumulation of AQP4 on astrocyte cell surfaces during osmotic or hypothermic challenge could be blocked with chelation of Ca^2+^ or antagonism of CaM, showing that Ca^2+^-CaM binding is essential for this trafficking process. Furthermore, phosphorylation of S276 in the C-terminal cytoplasmic domain was necessary but not sufficient for AQP4 translocation in response to hypotonicity [23].

The Ca^2+^-dependent binding of CaM to AQP4 was shown to be dependent upon core hydrophobic residues in a putative CBD motif located within the C-terminus of AQP4 [24]. While CaM-binding elicits a specific conformational change in the C-terminus of AQP4 and apparently controls cell-surface localization of the water channel, there is no precise structural description of CaM-binding events for this member of the AQP family. To further understand the molecular details involved in the regulation of AQP4 by CaM, we undertook a thorough characterization of the CaM-binding properties of AQP4. Herein, we demonstrate that AQP4 possesses Ca^2+^-dependent CBDs at both its cytosolic N- and C-termini. Additional complexity in AQP4 binding by CaM was revealed as the two CBDs in AQP4 display different modes for CaM interactions and could, in fact, bind simultaneously to the protein.

## 2. MATERIALS and METHODS

### 2.1. Materials

The human AQP4nt (Ac-^6^TARRWGK<u>S</u>GPL<u>S</u>TRENIMVAFKGVWT^31^-NH_2_; note the two Cys residues were modified to Ser (underlined) to reflect the reducing conditions in the cytoplasm of the cell) and AQP4ct (Ac-^256^VEFKRRFKEAFSKAAQQTKG SYMEV^280^-NH_2_) peptides were synthesized by Genscript Inc. (Piscataway, NJ, USA). Peptide sequences were confirmed by mass spectrometry and shown to be > 99% pure by analytical HPLC. Glutathione-Sepharose 4B, PreScission protease and pGEX-6P1 vector were purchased from GE Healthcare. All other chemicals were purchased from VWR Scientific or Sigma-Aldrich. Chicken calmodulin (CaM), which is identical in sequence to human CaM, was expressed in *Escherichia coli* and purified to homogeneity as described previously [26]. Uniformly ^15^N-labeled CaM was prepared in M9 minimal medium containing 0.5 g/L ^15^NH_4_Cl (Cambridge Isotope Laboratories) as the sole nitrogen source.

### 2.2. Expression and purification of recombinant rAQP4nt peptides

To identify experimentally the exact location of the CaM-binding domain in the N-terminal cytoplasmic region of human AQP4, we prepared a recombinant expression system for this region. Along with the recombinant rAQP4nt peptide, two variants including rAQP4ntW30A (P^6^TARRWGKSGPLSTRENIMVAFKGV<u>A</u>T^31^) and rAQP4nt6-19 (P^6^TARRWGKSGPLSTR^19^) were prepared. These peptides were expressed as insoluble ketosteroid isomerase (KSI) fusion proteins in LB medium. The His_6_-KSI-peptide fusion protein was then purified from inclusion bodies of *E. coli* using a Ni-column as previously described [27, 28]. The peptide was cleaved off at the Asp-Pro acid cleavage site that was implemented between the KSI protein and the peptide in 10% formic acid (80 °C for 90 min), leaving an extra Pro residue at the N-terminus of peptide as an artefact. After lyophilization, the released peptide could be resolubilized in buffer (20 mM Tris, 100 mM NaCl, pH 7.5), and it was further purified using a RP-HPLC with a Cosmosil Protein-R column (Nakarai Tesque Inc.) and lyophilized.

### 2.3. Expression and purification of recombinant rAQP4ct protein

A fragment of human AQP4 encoding the complete cytosolic C-terminal domain (rAQP4ct: residues 254 – 323; NP_001641.1) was amplified by standard PCR techniques and subcloned into the pGEX-6P1 vector. The sequence of the construct was verified by DNA sequencing. The GST-rAQP4ct fusion protein was produced in *E. coli* strain BL21(DE3) in LB medium. The fusion protein was isolated using glutathione-Sepharose 4B resin and cleaved on-column by treatment with PreScission Protease. The eluted rAQP4ct protein contained the cloning artifact ‘GPLGS’ at its N-terminus. The rAQP4ct protein domain was concentrated and exchanged into buffers for biophysical analyses with an Amicon centrifugal filter (Millipore).

### 2.4. NMR Measurements

All NMR experiments were performed at 30 °C on a Bruker Avance 700 MHz NMR spectrometer. Synthetic AQP4ct and AQP4nt peptides or recombinant rAQP4ct protein and rAQP4nt peptides were titrated into samples containing uniformly ^15^N-labeled CaM in the buffer containing 20 mM Bis-Tris (pH 7.0), 100 mM KCl, 6 mM CaCl_2_, 0.03% (w/v) NaN_3_, and 0.5 mM 3-(Trimethylsilyl)-1-propanesulfonate (DSS) in 90% H_2_O/10% D_2_O. Chemical shift perturbations (CSPs) upon addition of peptide were monitored by ^1^H, ^15^N-heteronuclear single quantum coherence (HSQC) NMR spectra. In some cases, the AQP4ct peptide was titrated into CaM complexed with AQP4nt to reach 1:1:1 molar ratios. The assignments of the ^1^H, ^15^N-HSQC signals of CaM were obtained as described previously [29]. All NMR spectra were processed with NMRPipe [30] and analysed using NMRView [31]. CSP values were calculated as a weighted average chemical shift difference of ^1^H and ^15^N resonances, using the equation CSP = (ΔHN)^2^ + (ΔN/5)^2^. Chemical shifts in all spectra were referenced using DSS [32]. In some HSQC titration experiments, dissociation constants (K_d_) were derived for the CBD peptide binding by curve fitting CSPs using several well-resolved peaks of CaM. The curve fittings were achieved using XCRVFIT v5.0.3 (developed by R. Boyko and B. D. Sykes, University of Alberta, Canada). The peptide and protein concentrations are measured using their predicted extinction coefficients at 280 nm (11000, 1490, and 2980 cm^-1^ M^-1^ for AQP4nt peptide, AQP4ct peptides and rAQP4ct construct, and CaM, respectively).

### 2.5. Isothermal Titration Calorimetry (ITC) Measurements

ITC experiments were performed on a MicroCal VP-ITC microcalorimeter at 30 °C. 0.26 mM of CaM in 20 mM HEPES (pH 7.5), 100 mM KCl and 1 mM CaCl_2_ were sequentially injected into a sample cell containing 28 μM of the recombinant rAQP4ct protein domain or the two synthetic peptides in the same buffer. Only the binding of the rAQP4ct domain produced useful ITC data. The data could be fitted to one set of sites-binding model using the MicroCal Origin software to obtain the K_d_ value.

### 2.6. Fluorescence Spectroscopy Measurements

Binding of the rAQP4nt and rAQP4ntW30A mutant to CaM were studied by exciting at 295 nm and recording the emission spectra between 300 and 450 nm. 10 uM peptide samples were prepared in the buffer containing 10 mM Tris, 100 mM KCl, and 2 mM CaCl_2_ at pH 7.5. Depending on the experiment, 12 μM CaM and 36 μM rAQP4ct protein were also added. The intrinsic Trp fluorescence was measured at room temperature on a Varian Cary Eclipse spectrofluorimeter. These intrinsic fluorescence experiments are facilitated by the fact that CaM and the rAQP4ct proteins do not contain any Trp residues.

## 3. RESULTS and DISCUSSION

### 3.1. Sequence analysis and comparison

In order to address the previously described impact of CaM antagonists on the regulation of AQP4 in brain and spinal cord cell swelling [23-25], we initiated a detailed biophysical investigation of the CaM-binding properties of AQP4. It is noteworthy that the cytosolic N-terminal (NT) and C-terminal (CT) sequences for all the human AQPs are poorly conserved within the family (Figure 1). Thus, we completed *in silico* assessments of the human NT and CT AQP4 sequences to identify any putative CBDs [8]. This inspection revealed two potential CBDs in AQP4: a novel one located in the NT sequence (Figure 2A), as well as the previously reported motif that is located in the CT domain sequence [24] (Figure 2B). In the process of examining AQP4 with the CBD predictor software, it appears that other human AQPs may also possess CBD motifs (Figure S1), although these have yet to be empirically validated and are not the specific subject of this report. It is noteworthy that AQP4 and AQP6 are amongst the isoforms in the AQP family that have an extended cytoplasmic NT domain. This might have important ramifications for AQP4 because the NT region of AQP6 was previously shown to contain a Ca^2+^-CaM-binding site [22], and AQP6 lacks a predicted CaM-binding site in the C-terminal domain. Moreover, AQP0 was demonstrated to possess a CBD within its cytoplasmic CT domain region [16-20], and the previously reported CBD in the CT of AQP4 possesses some gross similarity to that of AQP0 [24]. Additional *in silico* PSIPRED assessments of the secondary structure for the cytosolic NT and CT regions of AQP4 reveal that both can possess α-helical character that overlaps with the putative CBDs (bioinf.cs.ucl.ac.uk/psipred) [33]. Thus, herein the two CBD peptide sequences of AQP4 were displayed as helical wheels (Figure 2C) and also visualized as 3D helices (Figure 2D). Similar to the CBDs that are present in the NT of AQP6 and the CT of AQP0, the putative helical models of CBDs identified in NT and CT regions of AQP4 also describe amphipathic characteristics with hydrophobic and basic surfaces.

**Figure 1.**
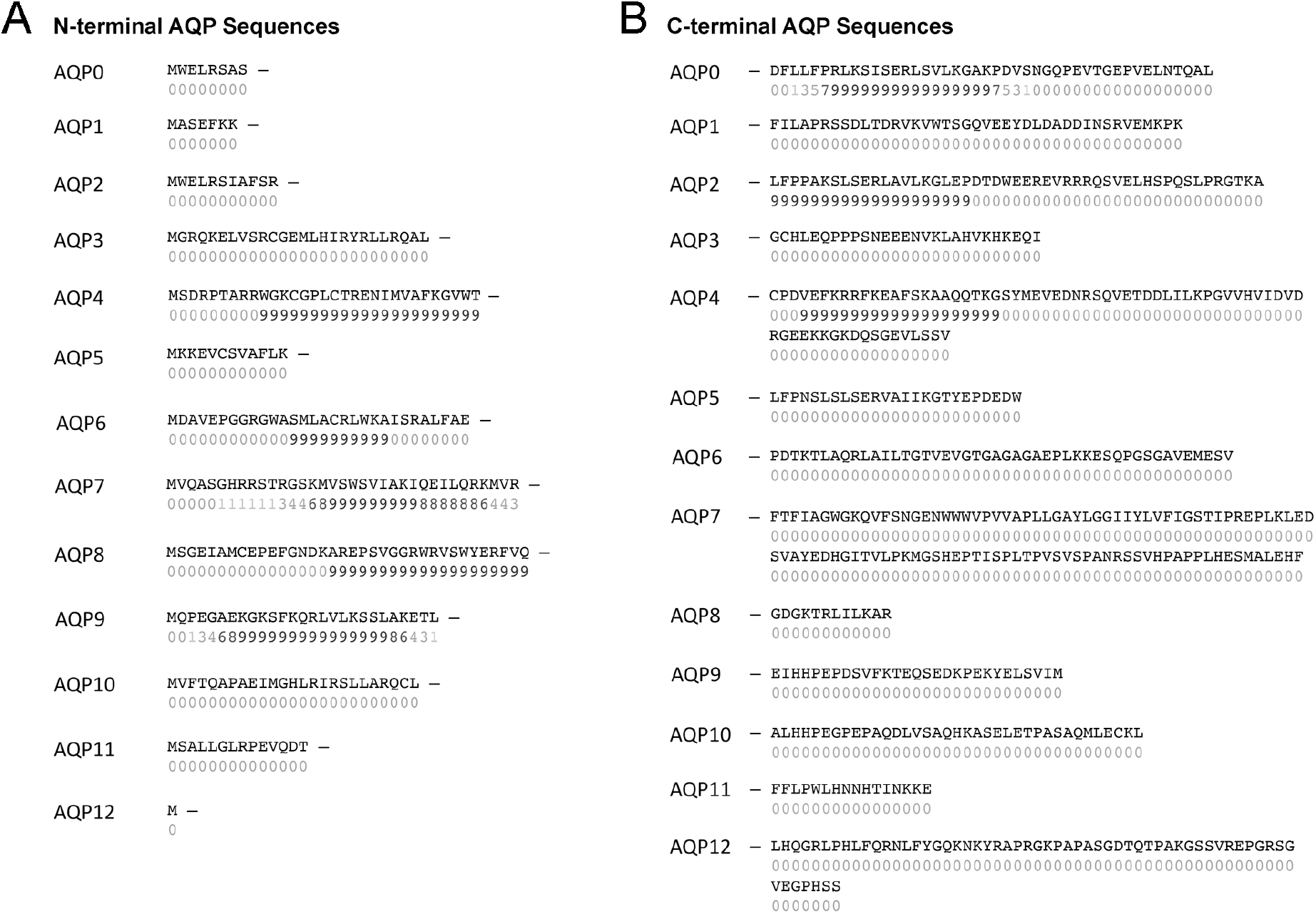
Location of predicted calmodulin-binding domains (CBD) in human aquaporin protein sequences. Amino acid sequences are provided for the cytoplasmic N-termini **(A)** and C-termini **(B)** of human aquaporin (AQP) proteins used in bioinformatic analyses to predict calmodulin-binding domains. The “Binding Site Search” function of the annotated Calmodulin Target Database was used. The sequences were evaluated based on primary structure (residue weight, charge, occurrence of particular residues including hydrophobic residues) as well as secondary structure (hydropathy, α-helical propensity, helical class) in a 20-residue sliding window. Scores (0-9) are given, and a string of high values (highlighted in dark blue) indicate the location of putative CaM-binding sites.

**Figure 2.**
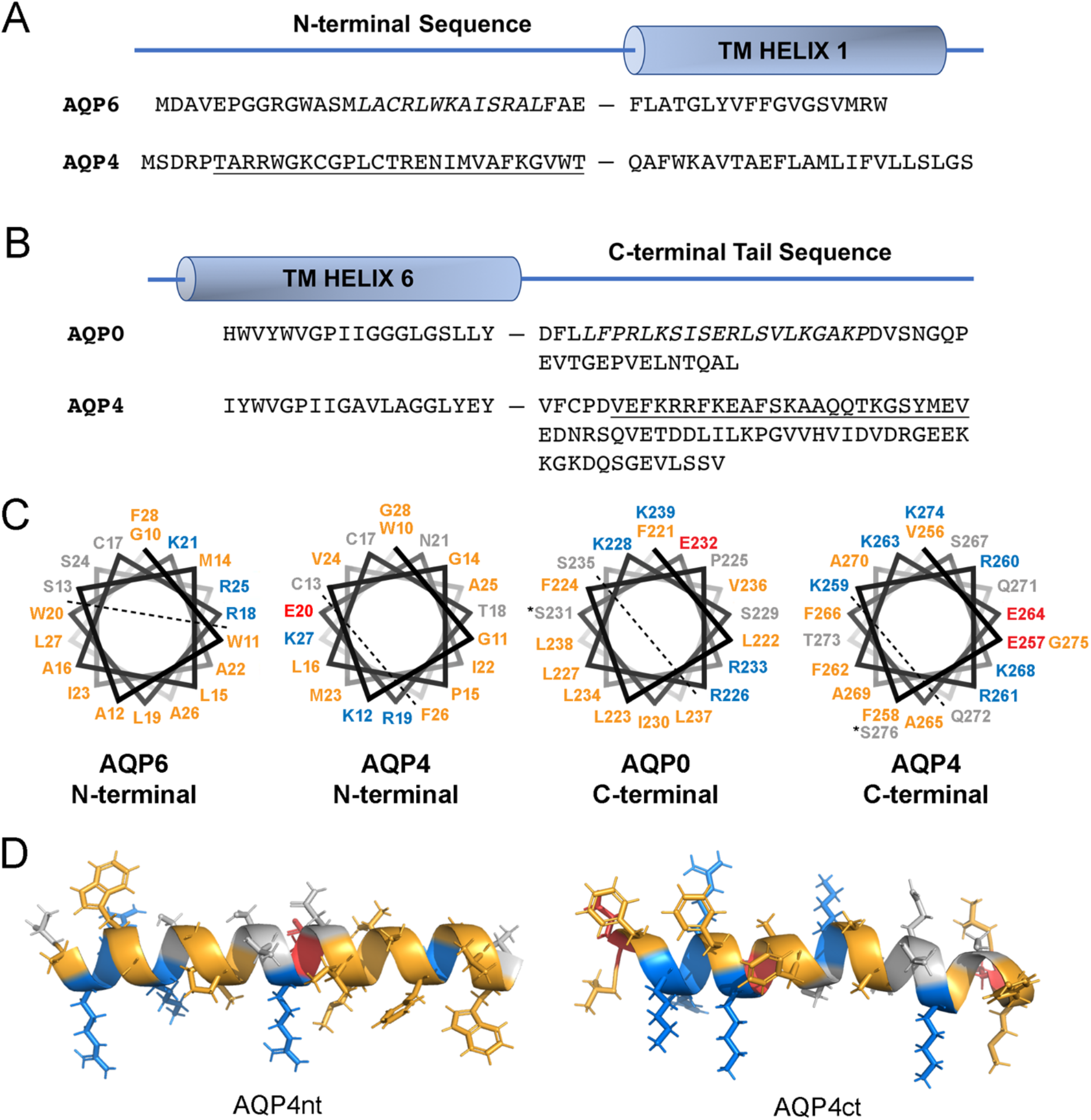
Analysis of the human AQP4 protein sequence identifies two putative CaM-binding domains. **(A)** An alignment of N-terminal cytoplasmic sequences of human AQP6 (NP_001643.2) and human AQP4 (NP_001641.1) preceding the first transmembrane (TM) helix. **(B)** An alignment of C-terminal cytoplasmic tail sequences of human AQP0 (NP_036196.1) and human AQP4 occurring after the final TM helix. The CBDs previously identified in AQP6 and AQP0 are indicated in italics. Putative CBD peptides (underlined) were synthesized and used to define CaM-binding properties. **(C)** Helical wheel projections of CBD motifs were generated for the AQP0, AQP6 as well as the AQP4ct and AQP4nt peptides. Dashed lines separate proposed hydrophobic and hydrophilic faces. Hydrophobic, acidic and basic residues are coloured yellow, red and blue respectively. *-indicates phosphorylatable residue. **(D)** 3D models of AQP4nt and AQP4ct peptides used in binding studies. Cys residues were replaced with Ser in the synthesized AQP4nt peptide. Colouring is as described previously.

The properties of the N-terminal CBD of AQP4 share reasonable similarity with the N-terminal CBD described for AQP6. Indeed, the CBD of AQP6 yielded an apparent K_d_ of 1.8 μM in titration experiments with dansylated CaM using fluorescence spectroscopy [22]. Although a complete structure determination is still pending, the AQP6 CBD site was putatively characterized as a 1-5-8-14 canonical binding motif. Additional hydrophobic residues located adjacent to these positions presumably contributed to CaM-binding since total abrogation of the interaction could only be elicited with replacement of several hydrophobic residues spanning the entire length of the sequence motif [22]. However, it should be noted that the K_d_ value indicates substantially weaker binding than typical canonical peptides, which often bind with a K_d_ in the nM range [2].

Previous studies with a 20-residue CBD peptide from the CT of bovine AQP0 describe a two-state binding model of interaction by ITC [19]. In this case, the step-wise binding revealed an initial high-affinity association (K_d1_: 0.07 μM) and a second lower-affinity event (K_d2_: 13 μM). Later, much weaker affinity values have been reported for the full-length AQP0 protein using microscale thermophoresis (MST) (K_d_: 2.5 and 40 μM) [34]. On the other hand, a single K_d_ value was previously determined by MST for the full-length, recombinant AQP4 protein solubilized in a detergent (K_d_ ∼30 μM) [24]. The CBD of AQP0 might mimic a 1-8-14 motif; the bovine CBD sequence of AQP0 contains hydrophobic anchoring residues at the three positions while human AQP0 lacks the third hydrophobic residue. The absence of a proper anchoring residue provides weaker CaM-binding affinity for human AQP0 when assessed with the dansyl-CaM fluorescence assay [16]. As expected, the lack of a hydrophobic anchoring residue in the human AQP4 CBD motif also appears to result in weaker CaM interaction [24]. While the CBD sequence present in the CT region of AQP4 shares some of the molecular characteristics of the CBD of AQP0, it also possesses a unique basic region upstream of the hydrophobic anchor. This sequence provides a basic amphipathic α-helix with one hydrophobic surface and one positively charged surface (Figure 2C). Much weaker affinities obtained with the AQP peptides compared to the canonical CBDs indicate that they are unlikely to follow a typical binding mode.

### 3.2. NMR and fluorescence spectroscopy studies with the rAQP4nt peptide

The ability of the AQP4nt and AQP4ct peptides to bind to Ca^2+^-CaM was assessed in detail with ^1^H,^15^N-HSQC NMR spectroscopy by titration assays with ^15^N-labeled CaM in which increasing amounts of the peptides were incubated with CaM in the presence of Ca^2+^. The AQP4nt peptide caused chemical shift perturbations (CSPs) in the spectra of ^15^N-labeled CaM, which appeared saturated at a 1:1 molar ratio and exhibited both fast and slow exchanging peaks on the NMR time scale (Figure 3). The majority of the fast and the slow exchanging peaks belong to the C- and N-lobe of CaM, respectively. Normally, slow exchange (K_ex_ < Δδ) represents a higher affinity binding event. Therefore, it appears that the AQP4nt peptide binds to CaM mainly through the N-lobe with a much weaker interaction with the C-lobe. To obtain an estimate of the affinity, some well-separated fast-exchanging peaks (e.g., G134, A102 and N137) were effectively fitted using a one-to-one binding model (Figure 3). However, the majority of the slow-exchanging peaks could not be fitted to a binding model due to severe peak broadening during the transition. Only I27 could be reasonably fit using the changes in the peak volume (Figure 3). These analyses determined an apparent K_d_ for this interaction to be 3.4 ± 0.8 uM (average value of four peaks). There were no data available to describe which lobe of CaM was involved in the interaction with AQP6 so a direct comparison with the binding mode of AQP4 is not possible at this time, but the determined K_d_ here is very similar to the reported Kd for CaM and the N-terminal CBD of AQP6 [22]. It is interesting that the AQP4nt peptide interacts preferentially with the N-lobe of CaM. Thus far, most peptides encompassing CaM-binding domains have shown stronger binding to the C-lobe of the protein [9]. However, there are a few published examples of peptides that display a preference for binding to the N-lobe of the protein [35, 36].

**Figure 3.**
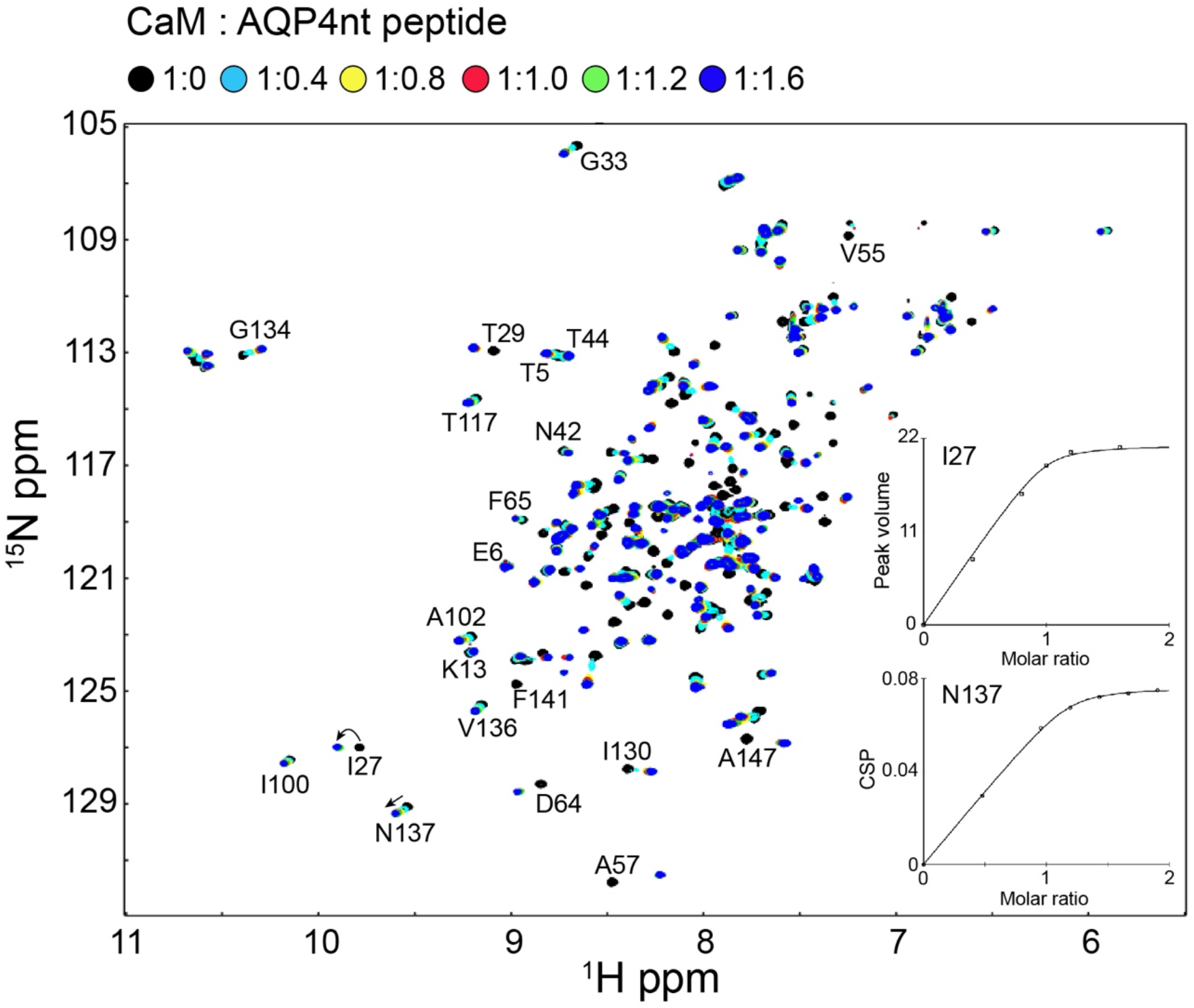
^1^H, ^15^N-HSQC-NMR spectroscopic analysis of CaM-AQP4nt peptide interactions. Overlaid ^1^H, ^15^N-HSQC-NMR spectra of 0.2 mM uniformly ^15^N-labeled CaM with increasing molar ratios of AQP4nt peptide (as indicated). Spectra were collected in Ca^2+^ buffer (20 mM Bis-Tris, 2 mM CaCl_2_, 100 mM KCl, pH 7.0). Peptide concentration-dependent chemical shift perturbations (CSPs) are plotted against AQP4nt concentration to determine K_d_ values for several residues (e.g., N137) that underwent fast exchange. Peak volumes were instead used for the residues undergoing slow exchange (e.g., I27).

For a better understanding of this unique binding, we used fluorescence spectroscopy. Trp residues that are present in the CBDs of target proteins generally serve as the anchoring residue for the C-lobe of CaM [9]. The N-terminal CBD of AQP4 contains two Trp residues (W10 and W30) (Figure 2). Figure 4A displays the steady-state Trp fluorescence of the AQP4nt peptide. The addition of CaM caused a significant enhancement of the intensity as well as a blue shift in the spectrum, meaning that the Trp residue is relocated to a more hydrophobic environment compared to the aqueous solution. Therefore, at least one of the two Trp residues in AQP4nt (W10 or W30) could act as an anchoring residue. We have therefore generated the W30A mutant of AQP4nt peptide. Since the AQP4nt-W30A peptide still caused a similar enhancement and a blue shift upon binding to CaM, we concluded that W10 is most likely the anchoring residue (Figure 4B). We then titrated the AQP4nt-W30A peptide into ^15^N-labeled CaM. The overlay of ^1^H, ^15^N-HSQC spectra showed very similar CSPs as the native peptide, suggesting that W30 is neither the anchoring residue nor critical for the binding (Figure 5A). As there are some changes observed mostly in the N-lobe signals, it seems that the mutation somewhat affects the N-lobe binding (Figure 5B). Thereby, the N-lobe could possibly interact with the C-terminal portion of the peptide near the mutation site. To further confirm this notion, we performed the same experiment with the AQP4nt-6-19 truncated peptide. As expected, this peptide only bound in fast exchange to the C-lobe of CaM (Figure 6). Hence, we conclude that CaM binds to the N-terminal CBD of AQP4 in an anti-parallel orientation: the C-lobe of CaM binds to the N-terminal portion of CBD with W10 as the anchoring residue, whereas the N-lobe of CaM binds to the C-terminal portion of CBD with an as yet unknown anchoring residue other than W30.

**Figure 4.**
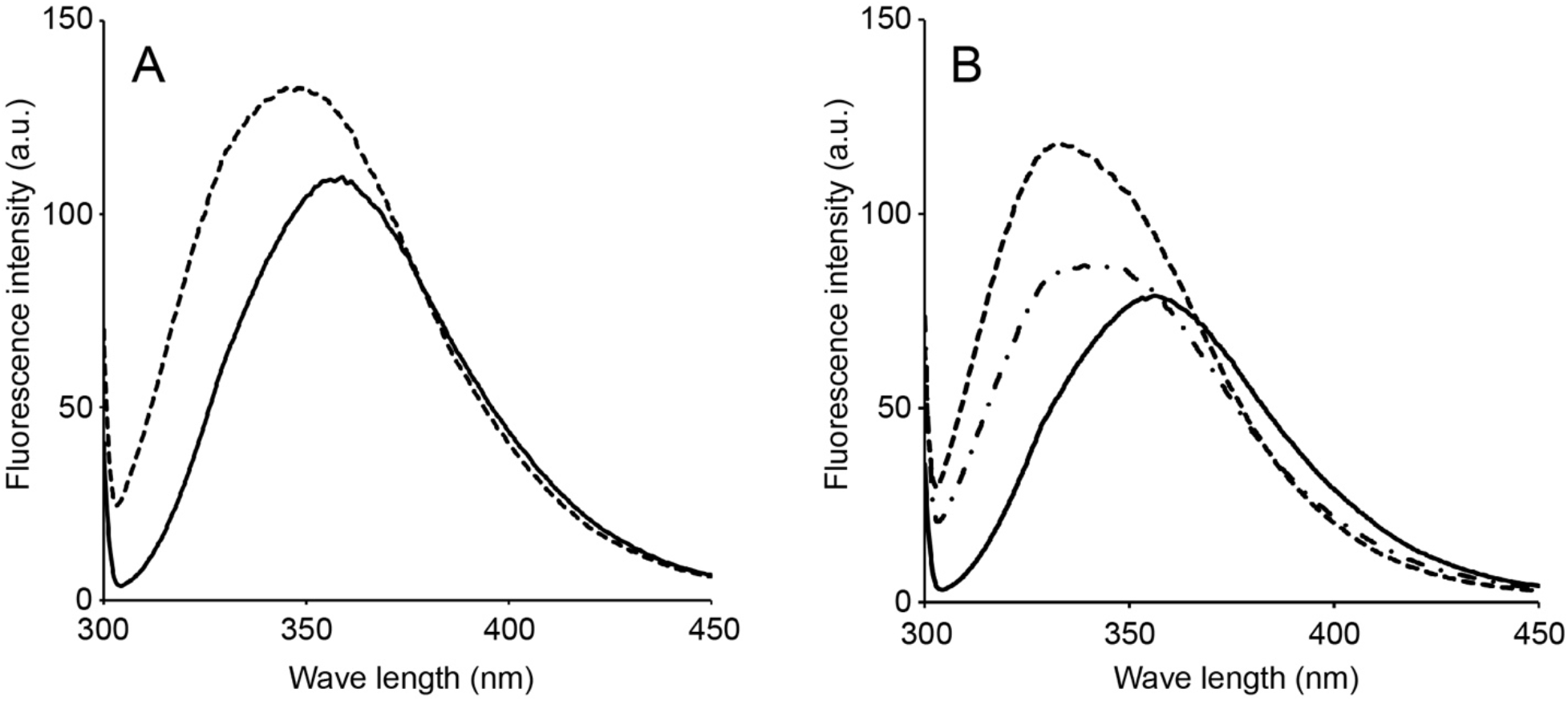
Steady-state Trp fluorescence studies of CaM-AQP4nt interaction. **(A)** Trp fluorescence of free (solid line) and CaM-bound AQP4nt peptide (dashed line). **(B)** Trp fluorescence of free (solid line) and CaM-bound AQP4nt-W30A (dashed line). rAQP4 construct was added to the same sample (dash-dotted line). Note that CaM and the rAQP4 construct do not possess any Trp residues.

**Figure 5.**
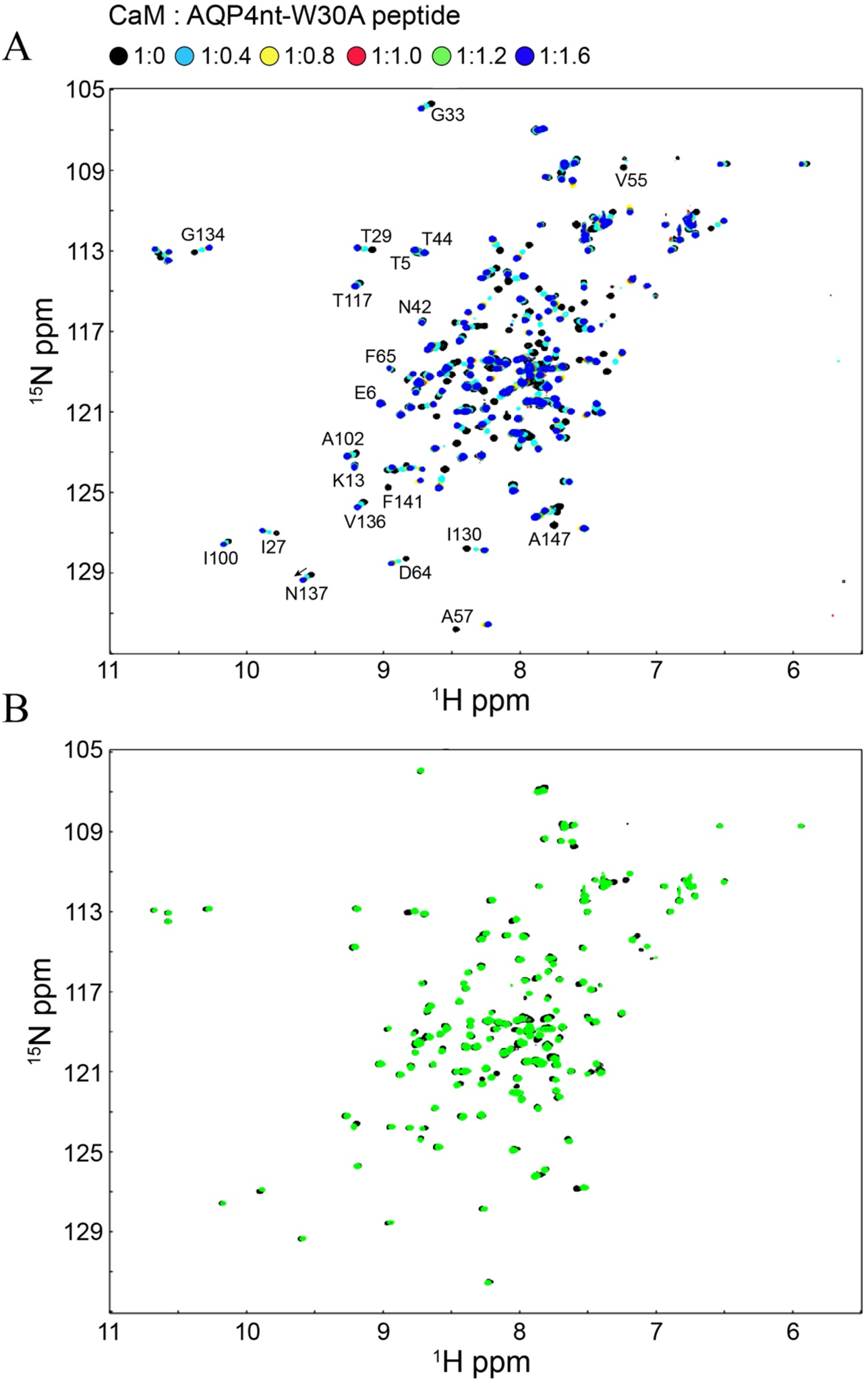
^1^H, ^15^N-HSQC-NMR spectroscopic analysis of CaM-AQP4nt-W30A mutant peptide interactions. **(A)** Overlaid ^1^H, ^15^N-HSQC-NMR spectra of 0.2 mM uniformly ^15^N-labeled CaM with increasing molar ratios of AQP4nt-W25A peptide (as indicated). Spectra were collected in Ca^2+^ buffer (20 mM Bis-Tris, 2 mM CaCl_2_, 100 mM KCl, pH 7.0). **(B)** Overlaid ^1^H, ^15^N-HSQC spectra of CaM-AQP4nt (black) and CaM-AQP4nt-W30A (green).

**Figure 6.**
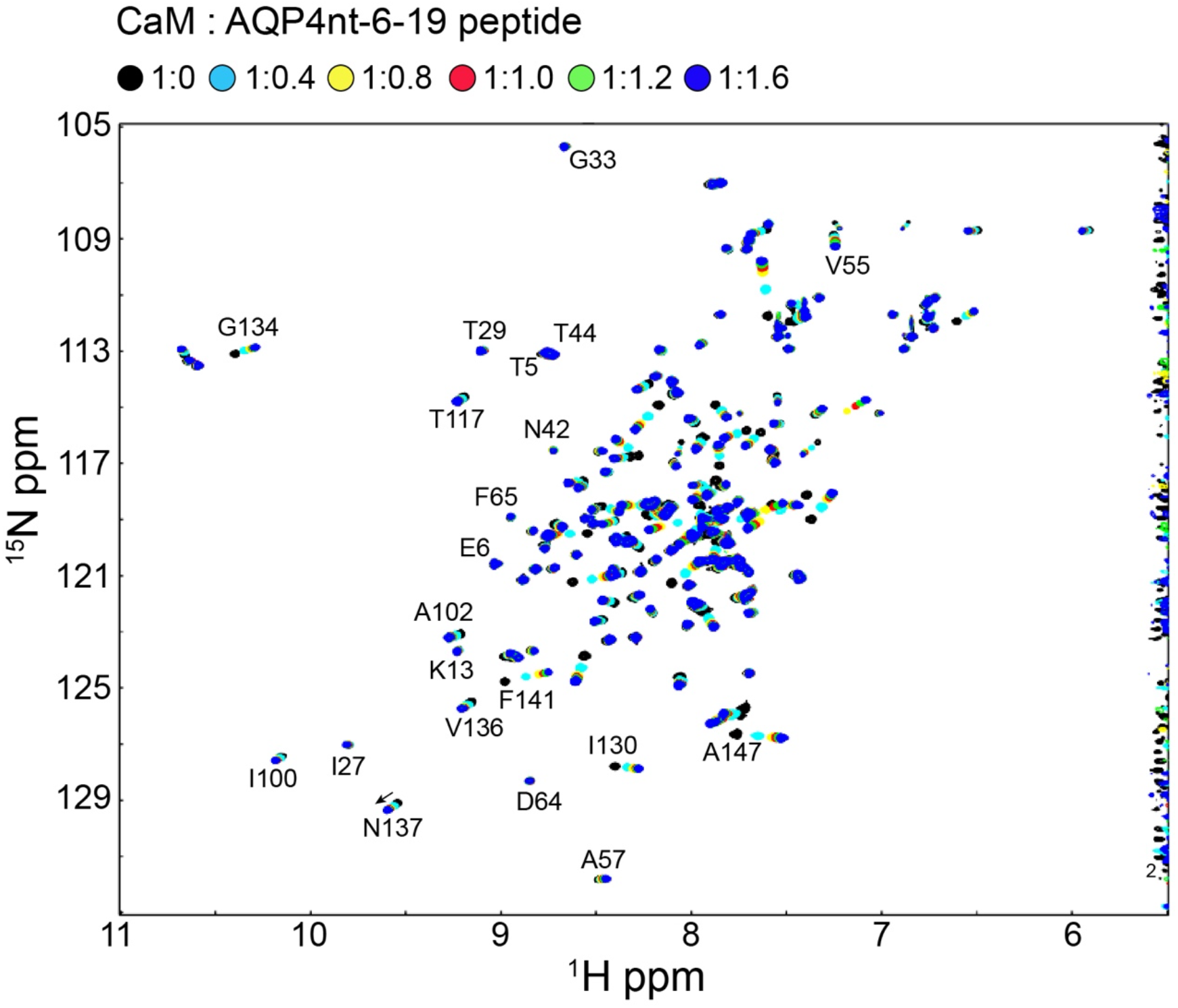
^1^H, ^15^N-HSQC-NMR spectroscopic analysis of CaM-AQP4nt-6-19 truncated peptide interactions. Overlaid ^1^H, ^15^N-HSQC-NMR spectra of 0.2 mM uniformly ^15^N-labeled CaM with increasing molar ratios of AQP4nt-6-19 peptide (as indicated in the figure). Spectra were collected in Ca^2+^ buffer (20 mM Bis-Tris, 2 mM CaCl_2_, 100 mM KCl, pH 7.0).

### 3.3. NMR studies of the synthetic AQP4ct peptide binding

Addition of the AQP4ct peptide also caused substantial chemical shift changes in the HSQC-NMR titration experiments (Figure 7A), exhibiting fast exchange on the NMR time scale in the spectra of ^15^N-labeled CaM. Fast exchange on the NMR timescale (Kex > Δδ) generally represents a relatively weak interaction (K_d_ 10 μM). The interaction between CaM and the AQP4ct peptide consisted of two distinct sequential binding events which appeared saturated at more than 1:2 molar ratio. The NMR data clearly demonstrate that the first binding event occurs on the C-lobe of CaM followed by a second binding event to the N-lobe (Figure 7B). From the CSPs mapped on to the ^15^N-labeled CaM structure, the AQP4ct peptide binds to the hydrophobic target binding pockets of CaM (Figure 7C). 55% of residues in the C-lobe with the most significant CSPs (> 0.1) are hydrophobic that includes I85, F92, L105, V108, M109, L116, V121, I125, I130, F141, V142, and M144. Similarly, 44% of residues in the N-lobe with the most significant CSPs (> 0.07) belong to hydrophobic residues that includes F12, F16, F19, V55, I63, L69, M71, and M76. The curve fittings of several well-separated peaks for the N-and C-lobe yielded apparent K_d_ values of 113.6 ± 8.0 and 3.6 ± 1.3 μM (average values of three peaks), respectively (Figure 7D). In contrast, the CaM-AQP0 interaction was mapped to both the N- and C-lobes with the dominant chemical shift changes also identified for residues located within the hydrophobic target-binding pockets of CaM [20].

**Figure 7.**
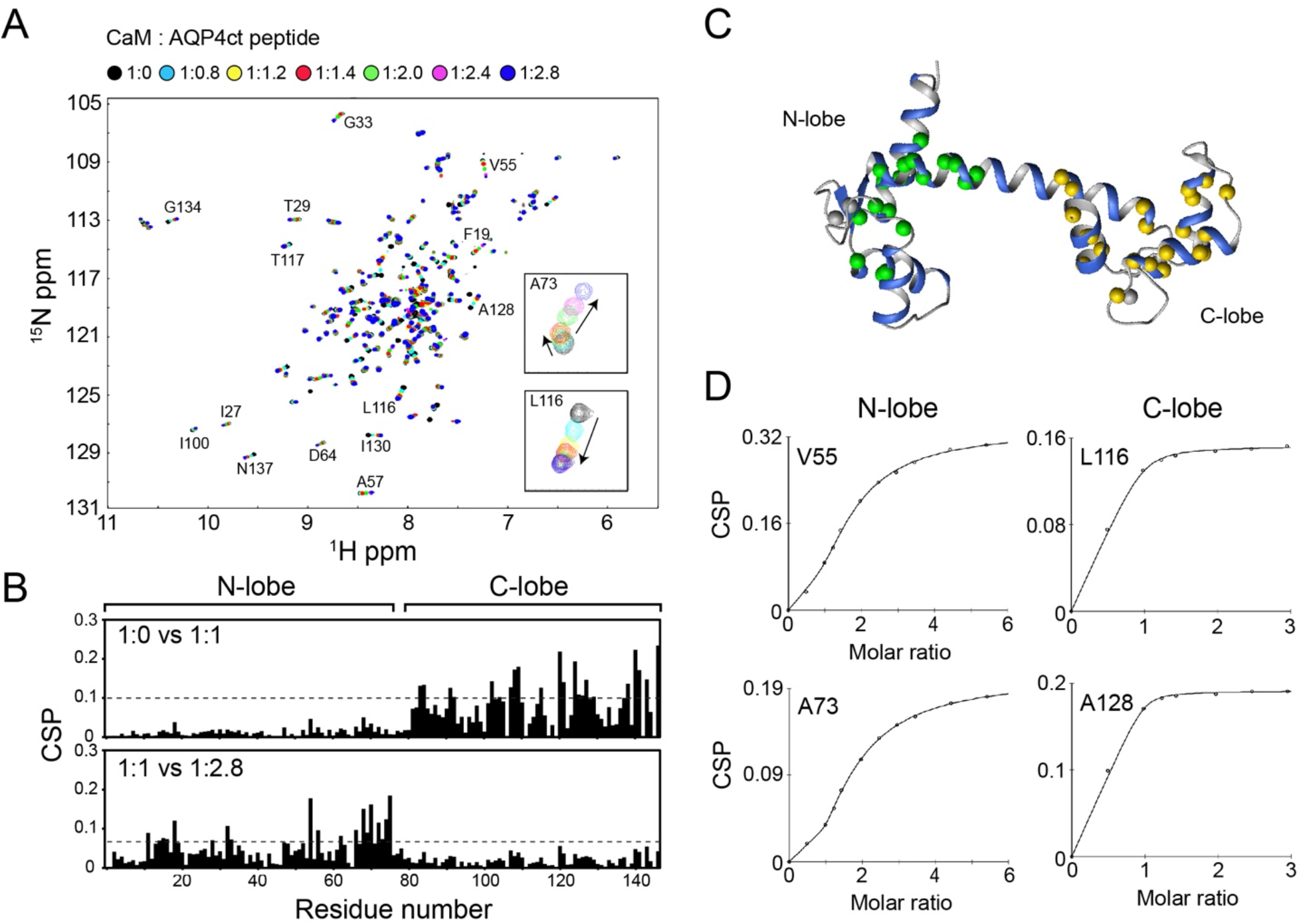
^1^H, ^15^N-HSQC-NMR spectroscopic analysis of CaM-AQP4ct peptide interactions. **(A)** Overlaid ^1^H, ^15^N-HSQC-NMR spectra of 0.2 mM uniformly ^15^N-labeled CaM with increasing molar ratios of AQP4ct peptide (as indicated). NMR samples were prepared in 20 mM Bis-Tris, 2 mM CaCl_2_, 100 mM KCl, pH 7.0. The peaks corresponding to two individual CaM residues, A73 and L116, are shown in enlarged panels. **(B)** The chemical shift perturbations (CSPs) induced by AQP4ct peptide binding to CaM are plotted as a function of the residue number. The location of the N- and C-terminal lobes of CaM are indicated. **(C)** The residues with a larger CSP (> 0.07 and 0.1 for N- and C-lobe, respectively) are highlighted on the Ca^2+^-CaM structure (1EXR). The residues that are involved in the first and the second peptide binding are coloured in yellow and green, respectively. **(D)** Examples of curve-fittings to derive K_d_ values for AQP4ct peptide bindings to the N- and C-lobe of CaM. While 1:1 binding mode was sufficient to fit the C-lobe binding, 1:2 sequential binding mode was required to fit the N-lobe binding.

### 3.4. Binding to apo-calmodulin

In Ca^2+^ and Na^+^ transmembrane channels, Ca^2+^-free CaM often associates via IQ motifs that are located in a cytoplasmic C-terminal domain [7, 37]. Such interactions are important as these are known to localize this regulatory protein and also modulate its affinity for Ca^2+^ [38, 39]. Therefore, we have also investigated the interaction between Ca^2+^-free CaM and the cytoplasmic C-terminal domain of AQP4. The addition of the rAQP4ct synthetic construct into ^15^N-labeled CaM in the presence of EDTA caused small but apparent chemical shift changes in ^1^H, ^15^N-HSQC spectra (Figure 8). The addition of AQP4ct peptide also caused very similar chemical shift changes (data not shown). Although amino acid sequence analysis did not identify any IQ or IQ-like motif in this region of the protein, there seems to be a weak association between CBD sequence and the C-lobe of CaM even in the absence of Ca^2+^. This weak interaction may contribute to transient localization of apo-CaM but is unlikely to affect Ca^2+^-binding.

**Figure 8.**
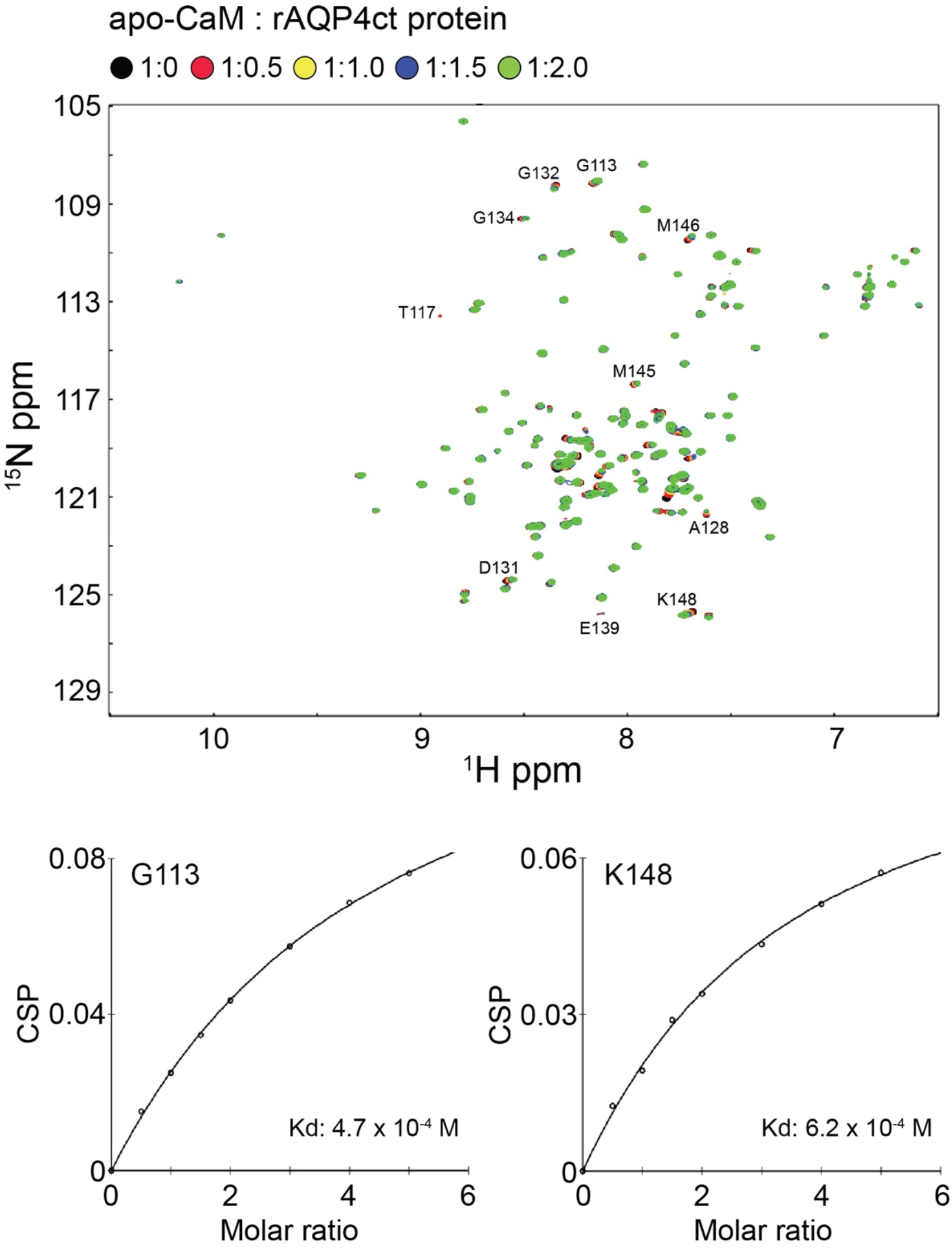
^1^H, ^15^N-HSQC-NMR spectroscopic analysis of apo CaM-rAQP4ct interactions. **(A)** Overlaid ^1^H, ^15^N-HSQC-NMR spectra of 0.2 mM uniformly ^15^N-labeled CaM with increasing molar ratios of rAQP4ct peptide (as indicated). Spectra were collected in EDTA buffer (20 mM Bis-Tris, 1 mM EDTA, 100 mM KCl, pH 7.0). **(B)** Peptide concentration-dependent chemical shift perturbations (CSPs) are plotted against rAQP4ct concentration to estimate K_d_ values for several residues (e.g., G113 and K148) that underwent fast exchange.

### 3.5. Binding of the entire C-terminal domain of AQP4

Given the non-canonical behavior of the CBD sequence motif in the CT region of AQP4, additional experiments were completed using a recombinant protein construct (rAQP4ct) that contained the entire cytosolic CT sequence. After purification of the rAQP4ct construct, CaM-binding properties were defined with ITC and NMR. Interestingly, the ITC data with rAQP4ct construct fit very well to a 1:1 binding model with the calculated affinity being a slightly weaker binding than for the AQP4ct peptide interaction with the C-lobe of CaM (Figure 9C; apparent K_d_ of 6.1 ± 3.3 μM). Titration of rAQP4ct with uniformly _15_N-labeled CaM and ^1^H,^15^N HSQC-NMR spectroscopy confirmed the 1:1 binding mode identified with ITC (Figure 9A). In this case, fast exchange was observed on the NMR timescale, which appeared saturated at a 1:1 molar ratio. Bound resonances were mapped for several residues in the C-lobe of CaM (Figure 9B and D). Although the residues involved in the binding are very similar to those observed with the AQP4ct peptide (∼95% identical), the magnitudes of the CSPs are somewhat smaller, which may suggest a slightly weaker affinity. Unlike the AQP4ct synthetic peptide, the CSPs observed in the N-lobe are almost negligible (Figure 9B). Therefore, we conclude that the rAQP4ct construct binds only to the C-lobe of CaM in a similar manner as the AQP4ct peptide. Although it is currently unclear why the rAQP4ct construct is unable to interact with the N-lobe of CaM, the regions downstream of CBD are likely to interfere with the N-lobe binding.

**Figure 9.**
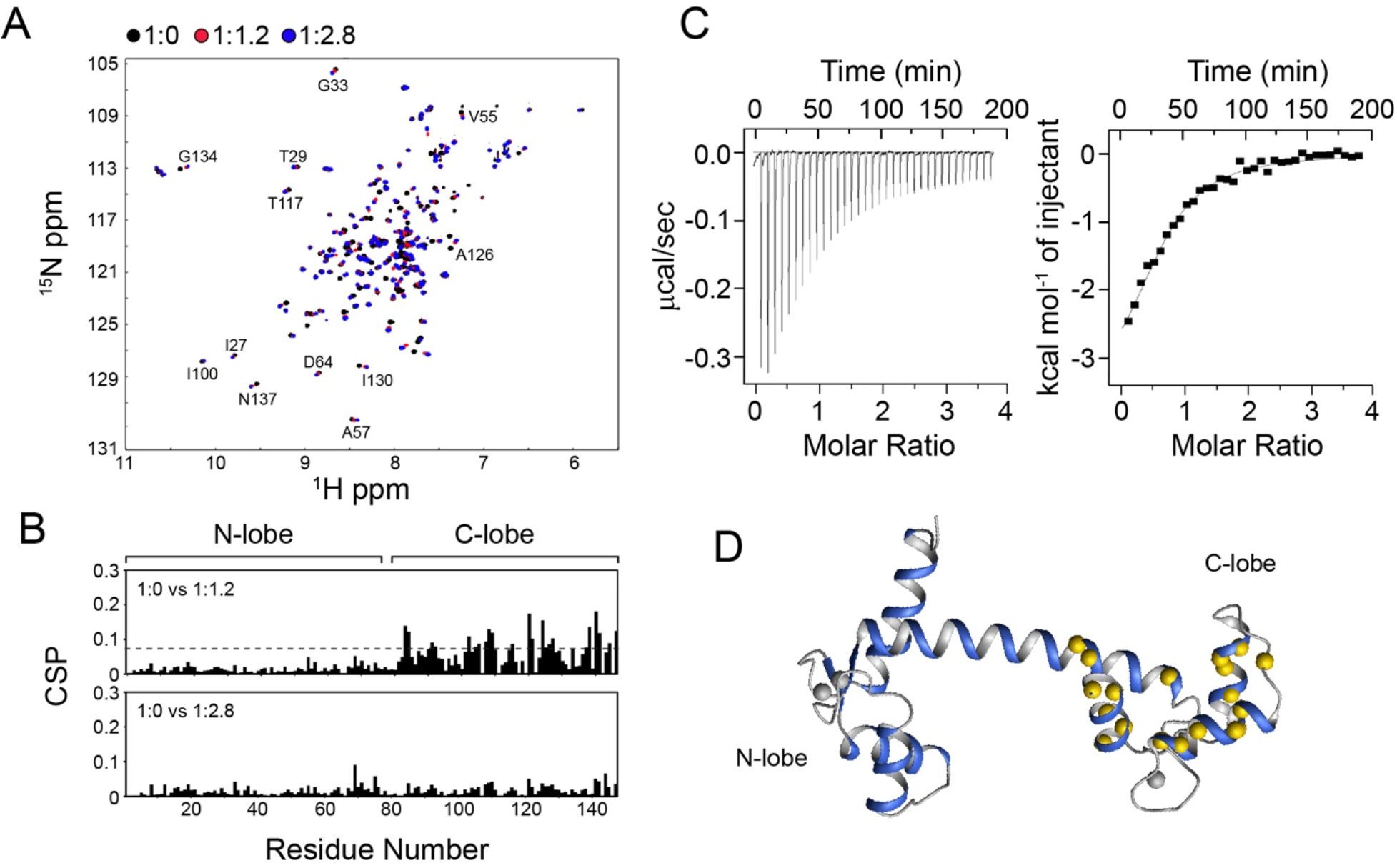
^1^H, ^15^N-HSQC-NMR spectroscopic analysis of CaM-rAQP4ct protein interactions. Overlaid ^1^H, ^15^N-HSQC-NMR spectra of uniformly ^15^N-labeled CaM in the presence of different molar ratios of recombinant rAQP4ct protein, 1:0 (black), 1:1.2 (red) and 1:2.8 (blue). The chemical shift perturbations (CSPs) induced by rAQP4ct protein binding to CaM are plotted as a function of the residue number. The location of the N- and C-terminal lobes of CaM are indicated. **(C)** Left, ITC raw heats of binding for Ca^2+^-CaM titrated into rAQP4ct protein. Right, binding isotherm with least-squares-fitted, one-site binding model. **(D)** The residues with a larger CSP (> 0.07) are highlighted on the Ca^2+^-CaM structure (1EXR).

### 3.6. Formation of a ternary complex

We further explored whether any higher order complexes could be generated with AQP4 and Ca^2+^-CaM since the CBD regions in both NT and CT regions of AQP4 demonstrated distinct binding interactions with preference for the N- and C-lobes of CaM, respectively. AQP4ct peptide was titrated into Ca^2+^–CaM that was previously complexed with AQP4nt peptide to yield a 1:1.1 ratio. Under these conditions, large chemical shift changes were observed mainly for the C-lobe of CaM while most of the peaks for the N-lobe remained almost identical (Figure 6). The results are suggestive of the formation of ternary complex with a 1:1:1 (CaM:AQP4nt:AQP4ct) stoichiometry. The pattern of chemical shift changes was somewhat distinct from that observed for CaM in the absence of AQP4nt peptide (Figure 10); thus, it is likely that association of AQP4nt peptide elicits some subtle structural impacts upon CaM prior to C-lobe occupancy by AQP4ct peptide. Moreover, the fitting of NMR titration data produced a K_d_ of 47.6 ± 2.4 μM (average of four peaks), which is ten times weaker than the binding affinity determined for the AQP4ct peptide interaction with the C-lobe of CaM in the absence of the AQP4nt peptide (Figure 10). We also observed the formation of the ternary complex using fluorescence spectroscopy. When the AQP4ct construct was added into the CaM bound AQP4nt-W30A peptide, the Trp fluorescence intensity was quenched, which indicates that W10 is re-exposed to the solvent as the C-lobe of CaM now engages with the rAQP4ct construct (Figure 4B).

**Figure 10.**
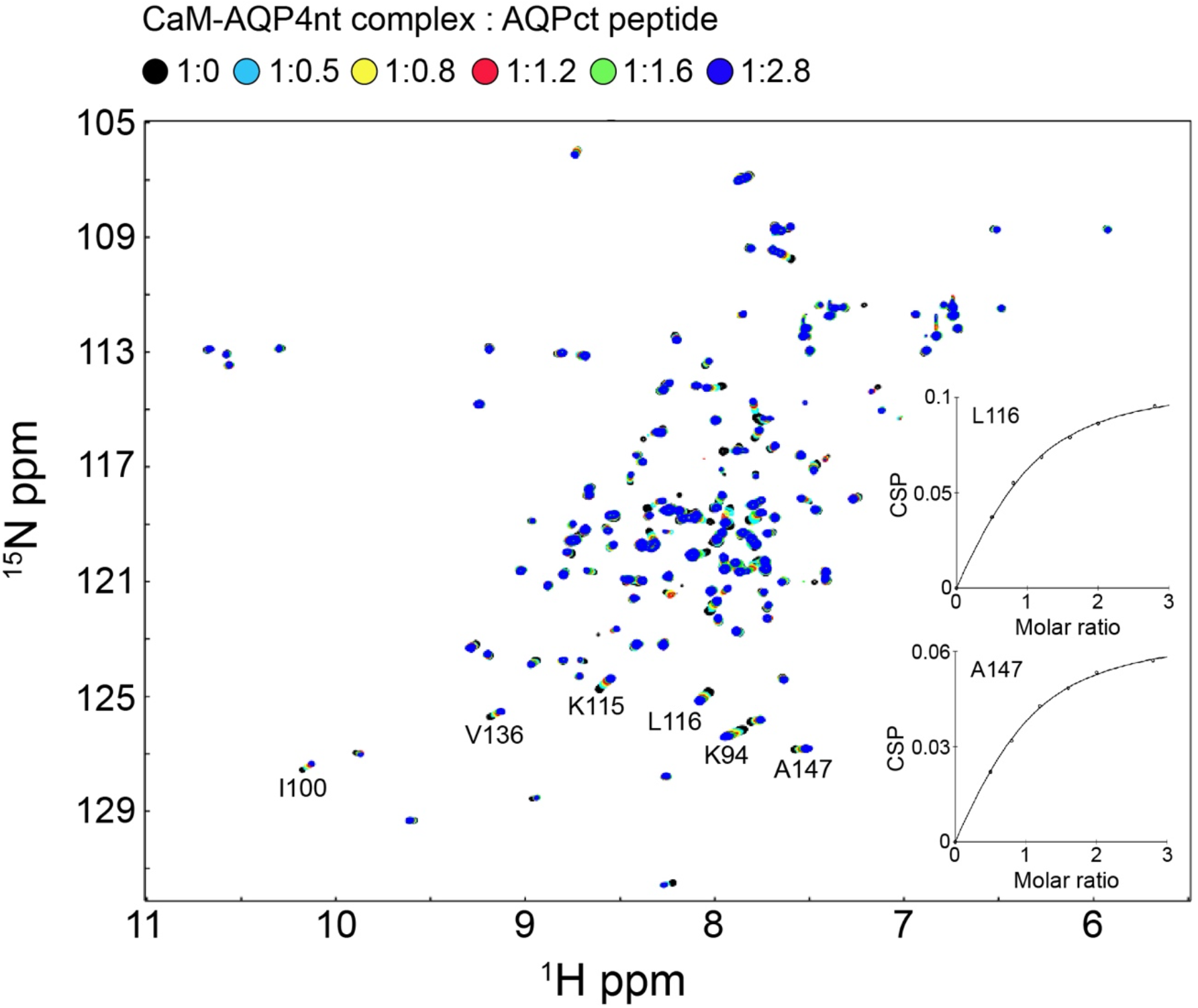
AQP4nt and AQP4ct peptides can simultaneously bind CaM. Overlaid ^1^H, ^15^N-HSQC-NMR spectra of 0.2 mM uniformly ^15^N-labeled CaM in the presence of AQP4nt peptide (in black) with subsequent titration of AQP4ct peptide at increasing molar ratios to give a final CaM:AQP4nt:AQP4ct ratio of 1:1.1:2.8 (in blue). Inset: The K_d_ value for binding of AQP4ct to the CaM:AQP4nt complex was determined from peptide concentration-dependent chemical shift perturbations (CSPs) obtained for several well-isolated peaks from the C-lobe of CaM (e.g. L116 and A147).

The potential functional importance of the CBD motif that we have identified here in the NT of human AQP4 is indicated by the observation that this region of the protein, like the C-terminal CBD of AQP4, is well conserved amongst diverse species (Figure 11). However, it is complicated by the fact that other studies indicate a complex and uncommon mechanism of translational control of AQP4 which permits the protein synthesis to occur from two different initiating methionines, M1 or M23 [40-42]. These two AQP4 isoforms are organized in the plasma membrane as heterotetramers. M23 is further observed to aggregate into structures known as orthogonal arrays of particles (OAPs) [41, 42], with the ratio of M23 and M1 determining the size of the OAP. The smaller M23 translation product of *AQP4* would likely not possess dual CaM-binding potential since most of the N-terminal CBD would be eliminated. Therefore, although it displayed the highest affinity for CaM *in vitro* of the two CBDs identified in AQP4, it is unclear what the ultimate biological impact of CaM-binding to the NT region of AQP4 would be. The M23 isoform appears dominant in the majority of tissues [42], but the inclusion of the NT and its inherent CaM-binding property could potentially contribute to the dynamic regulation of AQP4 during osmotic stress by influencing cellular localization. Furthermore, the two cysteine residues present in the NT CBD were reported to be palmitoylated in cultured cells [43], although it is not clear what proportion of the total M1 pool is palmitoylated, whether this differs between cell types, or to what extent this modification can be dynamically regulated. If this palmitoylation alters CaM binding affinity as would be expected, this may fine-tune the size of the CaM-accessible pool of M1.

**Figure 11.**
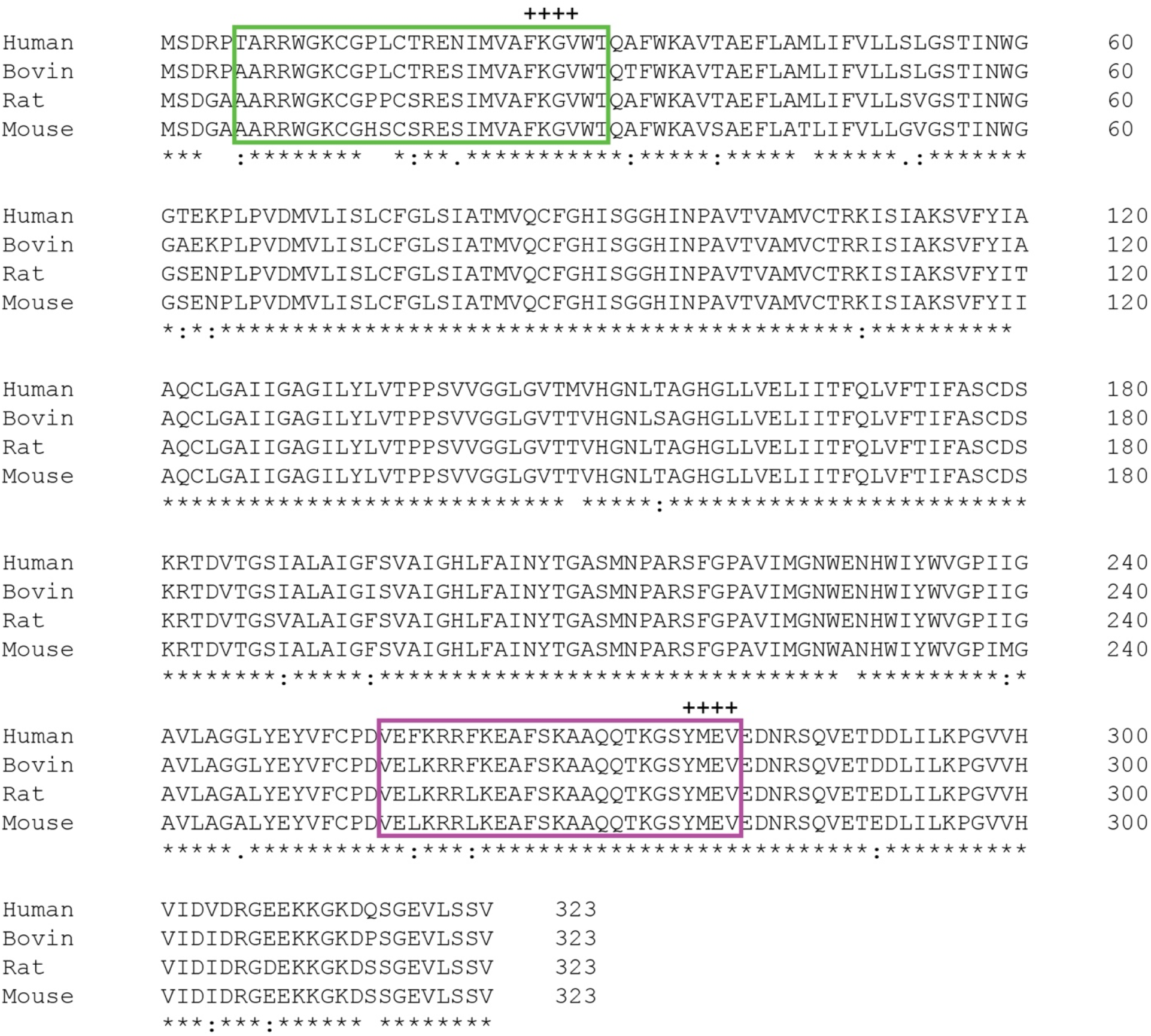
Sequence alignment shows that both the N- and C-terminal CBD regions are well conserved. The + symbols above the diagram indicate the location of the two separate internalization and trafficking motifs.

Finally, two internalization motifs (FXXV and YXXV; corresponding to N-terminal FKGV and C-terminal YMEV sequences in AQP4) are suggested to promote internalization and trafficking of various cell surface and transmembrane proteins [44], including AQP4 [41]. The FXXV motif is located within the N-terminal CBD, and the YXXV motif begins at the final hydrophobic anchoring residue of the C-terminal CBD of AQP4. Binding of CaM to these two CBD’s could therefore block access by other proteins to these interaction motifs. Thus, integration of CaM-binding potential adjacent to these sequences will likely be important for the regulation of AQP4 targeting. AQP4 has been suggested to form macromolecular complexes with a variety of ion channels, including the transient receptor potential (TRP) family members TRPV4 [45] and TRPM4 [46]. Both TRPV4 and TRPM4 contain CBDs [47, 48], so simultaneous binding of CaM to these channels and AQP4 via both the N- and C-lobes, may be a novel mechanism by which the stability and activity of these complexes could be modulated.

### 3.7. Effects of phosphorylation of Ser276

We have also studied the impact of AQP4 phosphorylation on the CT region since structural and functional studies have revealed phosphorylation to be a ubiquitous mechanism for AQP regulation [49]. A number of phosphorylation events have been reported to occur within the CT domain of AQP4. Indeed, the phosphorylation of S276 [23, 24, 50-52], S285 [50, 52, 53], Y277 [51, 54] and T289 [52] have been detected with high- and low-throughput studies. Intriguingly, both AQP0 and AQP4 possess a phosphorylatable Ser residue within the hydrophobic surface of the CBD peptide (Figure 2C). Phosphorylation of the S231 residue in AQP0 is known to inhibit the binding of CaM by modifying the interaction interface, suggesting a temporal regulatory mechanism for AQP0-CaM complex formation [17, 18, 20]. The S276 residue of AQP4 was also shown through mutational analysis to be phosphorylated *in vivo* and was linked to Ca^2+^-CaM-dependent, reversible translocation of AQP4 to the cell surface during extracellular hypotonic challenge of astrocytes [23, 24]. Phosphomimetic substitution of S276 with Glu appeared to strengthen the interaction of the AQP4 protein with CaM (i.e., ∼2-fold higher binding affinity as measured *in vitro* by MST) [24].

However, our comparison of CaM-binding to the synthetic AQP4ct and phospho-AQP4ct peptides by NMR spectroscopy did not reveal any significant differences in their binding modalities or affinities (Figure 12). The HSQC spectra of CaM bound to these two peptides are virtually identical at all the titration points, which indicates that S276 is not directly involved in this interaction. This observation is consistent with CSP’s measured previously in ^1^H,^15^N HSQC spectra of the ^15^N-labeled C-terminal domain, which indicated that binding of CaM predominantly affects the residues 255-275 [24]. It is important to note that the lack of apparent impact of S276 phosphorylation in our studies may arise from differences in reagents since previous MST data were collected for full-length AQP4 protein solubilized in detergent. For instance, the phosphorylation may induce some conformational changes in the full-length protein that are favorable for CaM-binding. Such disagreements have also been reported for AQP0 [34]. Further structural investigations would be required to understand the regulatory mechanism by the phosphorylation.

**Figure 12.**
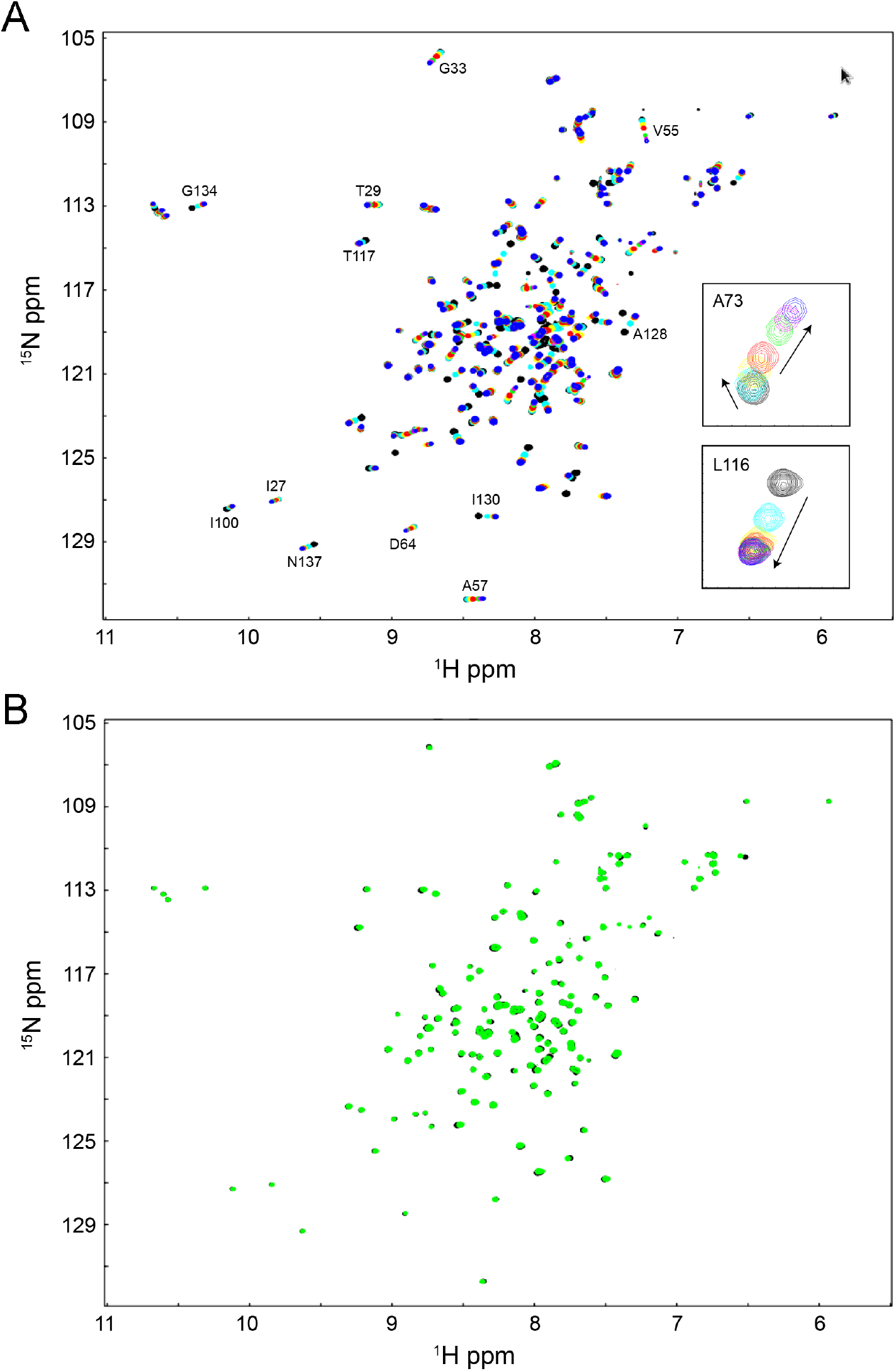
The interaction between Ca^2+^-CaM and phospho-AQP4ct peptide. **(A)** ^1^H, ^15^N-HSQC NMR spectra of ^15^N-labeled Ca^2+^-CaM with 0.0 (black), 0.8 (cyan), 1.2 (yellow), 1.4 (red), 2.0 (green), 2.4 (magenta), and 2.8 (blue) molar equivalents of phospho-AQP4ct peptide. **(B)** The HSQC spectra of Ca^2+^-CaM with AQP4ct peptide (black) and phospho-AQP4ct peptide (green) are overlaid at 1:2.8 protein/peptide ratios.

## 4. CONCLUSIONS

Taken together, our data support the notion of effective but complex molecular interactions between Ca^2+^-CaM and AQP4. The interactions between AQP4 CBDs and CaM determined in this study are summarized in Figure 13. The N-terminal CBD of AQP4 predominantly interacted with the N-lobe of CaM with a K_d_ of 10^−6^ M; this is in contrast to the C-terminal CBD which binds predominantly to the C-lobe of CaM with a similar affinity (K_d_ ∼10^−6^ M). Ultimately our data support a binding model whereby CaM can form a ternary bridging complex with the two AQP4 CBD’s (Figure 13), where Ca^2+^-CaM seems to act like an adapter protein, bridging between two parts of the protein. Our observations imply that unlike AQP0, CaM could bind to AQP4 with 1:1 stoichiometry in a parallel orientation, where the N- and C-lobes of CaM interact with the N- and C-terminal CBDs of AQP4, respectively. However, our results do not exclude the possibility for CaM to bridge between two distinct AQP4 molecules within a tetramer, or even between separate tetramers, Be that as it may, our results clearly highlight a difference between the Ca^2+^-CaM interactions of AQP0 and AQP4, suggesting that these different binding modes may contribute to the different biological roles of the two AQP isoforms, a predominantly structural role in the case of APQ0, versus the regulation of the trafficking from the storage vesicle to the plasma membrane in the case of AQP4. Given our results, additional analyses of the interdependency of protein-phosphorylation, -palmitoylation and CaM-binding on the vesicular trafficking of AQP4 as well as on the oligomerization and formation of OAPs for AQP4 are warranted.

**Figure 13.**
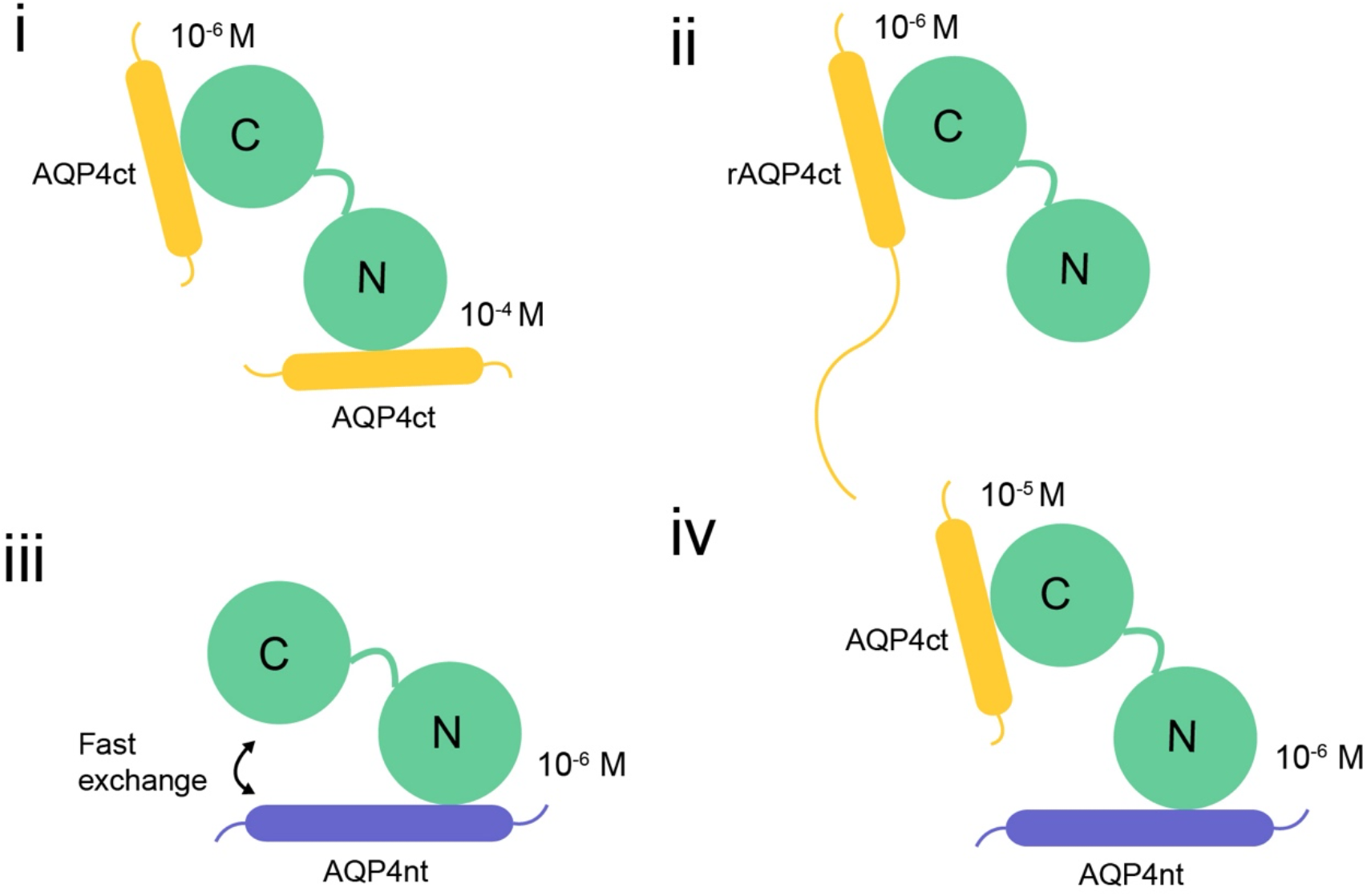
Binding models of AQP4-CBD peptides. (i), AQP4ct peptide (ii), recombinant rAQP4ct protein, (iii) AQP4nt peptide, and (iv) combined AQP4nt-AQP4ct peptides for Ca^2+^-CaM.

## 5. AUTHOR CONTRIBUTIONS

Hiroaki Ishida: Conceptualization, Investigation, Formal analysis, Visualization, Writing - Original draft preparation. Hans Vogel: Conceptualization, Supervision, Writing – Review and Editing, Funding Acquisition. Justin MacDonald: Conceptualization, Resources, Visualization, Writing - Original draft preparation, Funding Acquisition. Philip Kitchen: Resources, Writing – Review and Editing, Funding Acquisition. Roslyn Bill: Resources, Writing – Review and Editing, Funding Acquisition. Alex Conner: Resources, Writing – Review and Editing, Funding Acquisition.

## 6. ACKNOWLEDGEMENTS

This work was supported by grants from the Natural Sciences and Engineering Research Council (NSERC; RGPIN/04379-2019 to JAM, and RGPIN/05977-2015 to HJV) and the Biotechnology & Biosciences Research Council (BBSRC; BB/P025927/1 to ACC, PK and RMB). A BBSRC International Partnering Award (BB/P025027/1 to RMB and JAM) also facilitated research activities. The authors thank Mona Chappellaz for technical assistance in generating the rAQP4ct expression construct.

